# Live-cell assays for cell stress responses reveal new patterns of cell signaling caused by mutations in rhodopsin, α-synuclein and TDP-43

**DOI:** 10.1101/735878

**Authors:** Kevin M. Harlen, Elizabeth C. Roush, Joseph E. Clayton, Scott Martinka, Thomas E. Hughes

## Abstract

Many neurodegenerative diseases induce high levels of sustained cellular stress and alter a number of cellular processes. Genetically-encoded fluorescent biosensors are effective tools to examine neuronal activity and signaling in living cells. To examine how different mutations associated with neurodegenerative disease affect cell stress and signaling we created live-cell assays for ER-mediated cell stress and second messenger signaling. Analysis of the rhodopsin P23H mutation, the most common mutation in autosomal dominant Retinitis Pigmentosa, revealed increased cell stress levels compared to wild type rhodopsin. Moreover, this increase in cell stress correlated with blunted Ca^2+^ signaling in a stress dependent manner. Analysis of single cell Ca^2+^ signaling profiles revealed unique Ca^2+^ signaling responses exist in cells expressing wild type or P23H mutants, further supporting the notion that second messenger signaling is affected by cell stress. To explore the use of the ER-stress biosensor in other neurodegenerative diseases we examined how various mutants of α-synuclein and TDP-43 affected ER-mediated cell stress. Mutants of both α-synuclein and TDP-43 associated with Parkinson’s Disease and ALS demonstrated increases in ER-mediated cell stress. This increased cell stress was accompanied by changes in phosphodiesterase activity. Both HEK293T and SH-SY5Y cells expressing these proteins displayed a shift towards increased cAMP degradation rates, likely due to increased phosphodiesterase activity. Together these data illustrate how biosensors can provide nuanced, new views of neurodegenerative disease processes.

## INTRODUCTION

Genetically-encoded fluorescent biosensors are powerful tools that have provided new views of how circuits in the brain operate, and how cells and networks of cells process and respond to stimuli (Chen et al., 2017). Many of these biosensors detect small molecule analytes, like Ca^2+^, cyclic AMP (cAMP) and diacylglycerol (Broussard et al., 2014; Tewson et al., 2012, 2016; Zhao et al., 2011) that change in concentration during a cell signaling events. Biosensors of this type are used to monitor events such as G-protein coupled receptor (GPCR) activation or neuronal Ca^2+^ signaling. Because of their genetic nature, biosensors can be targeted to distinct sub-populations of cells or to specific organelles and subcellular domains (Pendin et al., 2017), and can be systemically delivered by viral vectors. These features have popularized genetically-encoded biosensors, especially those for Ca^2+^ and cAMP (Castro et al., 2014), for use in both in vitro as well as in vivo models (Chen et al., 2017). Another type of genetically encoded biosensor targets changes to the state of the cell. For example, biosensors for apoptosis (Xu et al., 1998), cell cycle state (Sakaue-Sawano et al., 2008), autophagy (Katayama et al., 2011), and cell stress (Iwawaki et al., 2004; Roy et al., 2017) have been developed to detect broad changes to cellular states. However, these two classes of biosensors are often used in separate assays to examine unique outcomes of either changes in cell state or signaling. We reasoned that combining biosensors for cell state with those for cell signaling could provide a means to assess how changes in cell state, such as cell stress, alter cellular signaling. An example where changes in cell state and signaling overlap is in neurodegenerative disease. Neurodegenerative disorders, such as Parkinson’s Disease, Amyotrophic lateral sclerosis (ALS), and the degenerative blinding disease Retinitis Pigmentosa (RP) all involve cellular stress and occur over the course of many years. Each disease is also linked to changes in second messenger signaling. In RP, rod photoreceptors slowly degrade over time, eventually degrading the cone photoreceptors as well, leading to blindness (Ferrari et al., 2011; Hartong et al., 2006). The most common mutation associated with RP is the autosomal dominant P23H mutation within the rhodopsin gene (Ferrari et al., 2011). This loss of function in rhodopsin is accompanied by changes in Ca^2+^ and cyclic GMP (cGMP) signaling, along with increased cell stress triggered by the unfolded protein response (UPR) (Arango-Gonzalez et al., 2014; Shinde et al., 2016). In Parkinson’s and ALS, unique subpopulations of neurons experience prolonged cell stress before eventually dying (Bosco et al., 2011; Maiti et al., 2017; Taylor et al., 2016). Parkinson’s and ALS are characterized by the accumulation of misfolded proteins throughout the cell. However there is also evidence that the modulation of second messenger signaling levels mediated through GPCR activity may influence disease progression (Mittal et al., 2017; Xu et al., 2012). Furthermore, inhibition of phosphodiesterase activity, which is responsible for the breakdown of cAMP and cGMP, has been demonstrated to preserve dopaminergic neurons in models of Parkinson’s disease (Morales-Garcia et al., 2011). Thus, accumulating evidence suggests that changes in both cell stress and signaling are associated with multiple neurodegenerative diseases.

Importantly, these changes in stress and signaling occur throughout the progression of the disease, while neurons are still alive and presumably functional. However, post mortem studies and animal models often focus on the cell death that ultimately occurs. But dead cells tell no tales. Imagine the neuron suffering under the load of a misfolded protein for years on end. Which stress pathways are activated and how does it compensate for the stress? Does it still respond to its environment appropriately? Can it still sense the same neurotransmitters and neuromodulators in the same way? Does it still respond to drugs in the same way that the healthy, surrounding cells do? To begin to assess these questions we created a biosensor to detect endoplasmic reticulum (ER)-mediated cell stress, which is associated with RP, Parkinson’s and ALS (Remondelli and Renna, 2017; Shinde et al., 2016), and paired it with biosensors for second messengers. We then monitored, either simultaneously or orthogonally, changes in cell stress levels and the effects on cellular signaling. Using mutations of rhodopsin, α-synuclein and TAR DNA binding protein (TDP-43) we show that Ca^2+^ and cAMP signaling is altered under cell stress.

## MATERIALS AND METHODS

### Biosensor construction

The ER-stress biosensor was modified from (Iwawaki et al., 2004). mNeonGreen was placed downstream of the XBP1 sequence and splice site. SantakaRFP (ATUM FPB-58-609) was placed upstream of the XBP1 sequence (Figure 1A). The SantakaRFP and XBP1-mNeonGreen sequences were separated by a self cleaving 2A peptide sequence. For the R-GECO-cell stress biosensor SantakaRFP was replaced with R-GECO1.2 (Wu et al., 2013) (Figure 3A). Cloning was done using In-Fusion HD cloning (Takara 638911). R-GECO (U0600R), R-cADDis (U0200R) and bPAC (V0100N) were obtained from Montana Molecular.

**Figure 1.**
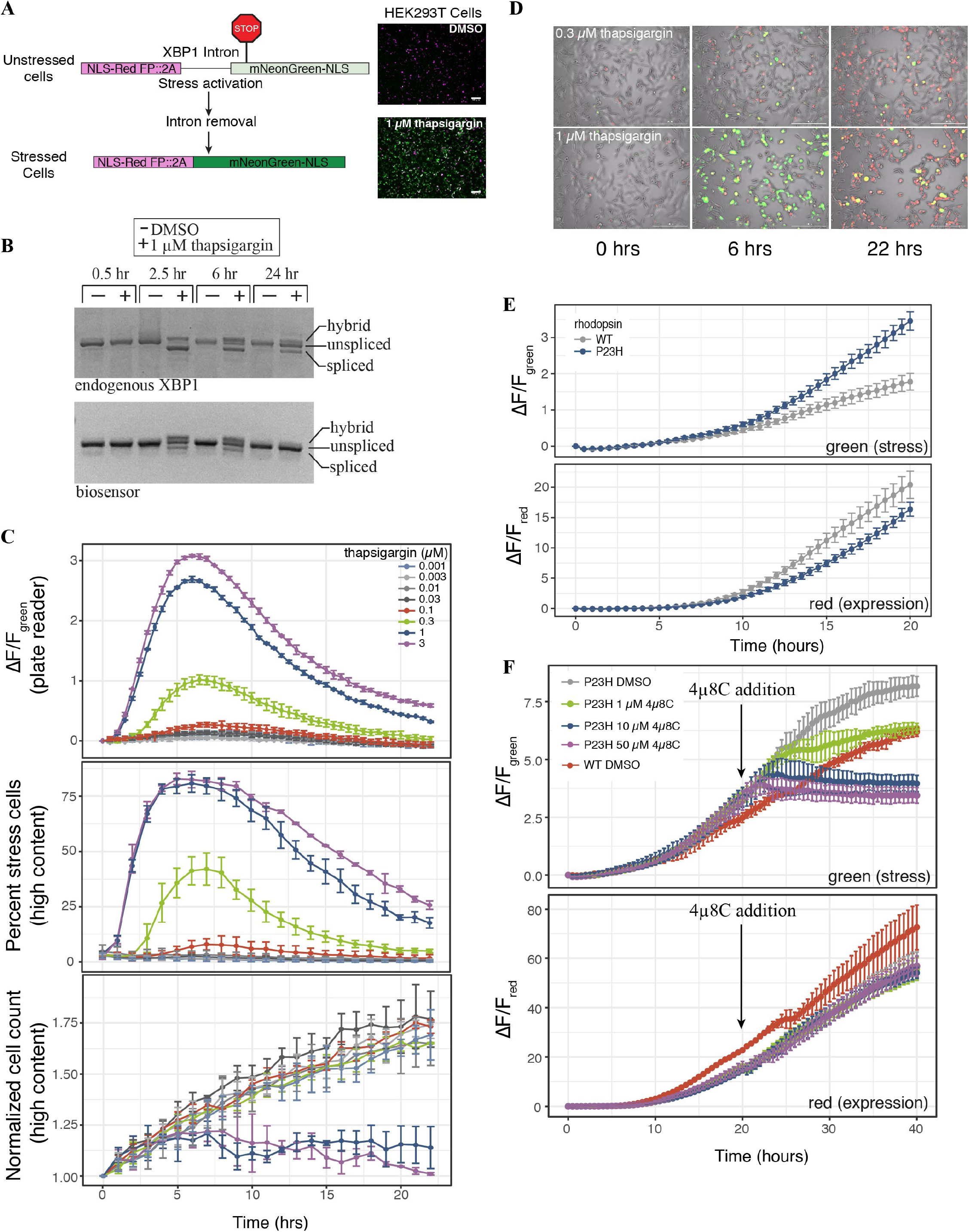
A genetically-encoded fluorescent biosensor to detect ER-mediated cell stress. **(A)** schematic of the two-color cell stress biosensor. The biosensor is polycistronic transcript consisting of a nuclear targeted constitutively expressed red fluorescent protein followed by a self cleaving 2A peptide and a stress-induced nuclear targeted green fluorescent protein (mNeonGreen) fused to a portion of the XBP1 protein. Activation of ER stress through the IRE1α-XBP1 pathway splices out a portion of the XBP1 mRNA, shifting the mNeonGreen into frame. Top: unstressed cells; bottom: stressed cells treated with 1 μM thapsigargin. Scale bar = 100 μm. **(B)** RT-PCR agarose gel of the splicing status of either endogenous or biosensor XBP1 transcripts over 24 hours. Cells were treated with either DMSO or 1 μM thapsigargin. Three bands are present, unspliced XBP1, spliced XBP1 and a hybrid band of spliced and unspliced transcript (Chalmers et al., 2017). **(C)** HEK293T cells were transduced with the cell stress biosensor and treated with increasing amounts of thapsigargin. Top: the fold change (ΔF/F) in green fluorescence was calculated over 22 hours, measured every 30 minutes on a plate reader. Middle: the percent of stressed cells per image was calculated every hour on a high content imager. Bottom: the normalized cell count per image was calculated every hour on a high content imager. Data are plotted as mean ± s.d., n = 3 wells per condition. **(D)** Overlay of phase, red fluorescence, and green fluorescence images of HEK293 cells transduced with the cell stress biosensor and treated with 0.3 μM and 1 μM thapsigargin. Images acquired at 0, 6 and 22 hours after treatment, scale bar = 200 μm. **(E)** HEK293T cells were transfected with the cell stress biosensor and either wild type (WT) or mutant P23H rhodopsin. The fold change in green (stress) and red (expression) fluorescence was measured over 20 hours. Data are plotted as mean ± s.d., n = 3 wells per condition. **(F)** After 20 hours the cells from (D) were treated with increasing amounts of the IRE1α inhibitor 4μ8C or DMSO. The fold change in green and red fluorescence was then monitored for an additional 20 hours.

### Chemicals

Sodium butyrate (B5887), valproic acid (P4543), and carbachol (C4382) were obtained from Millipore Sigma. Thapsigargin (10522) and 4μ8C (22110) were obtained from Cayman Chemical.

### Cell culture

HEK293T cells were obtained from ATCC (CRL-11268) and cultured in T-75 flasks at 37°C and 5% CO2 in EMEM media (ATCC 30-2003) containing 1× Penicillin-Strep-tomycin (Thermo Fisher 15140122), supplemented with 10% fetal bovine serum (FBS). SH-SY5Y cells were obtained from ATCC (CRL-2266) and cultured the same as the HEK293T cells except DMEM (30-2002) was used as the base meda. For plate reader experiments cells were cultured in Fluorobrite DMEM (Thermo Fisher A1896701) containing 4 mM Glutamax (35050061) and 10% FBS (Figure 1), or the media was replaced with DPBS (Figures 2-5). For imaging media was replaced with DPBS containing calcium and magnesium (Fisher Scientific SH30264.01). For all experiments cells were subcultured at 70-90% confluence and plated onto 96-well poly-D lysine coated microplates (Greiner Bio-One 82050-806).

**Figure 2.**
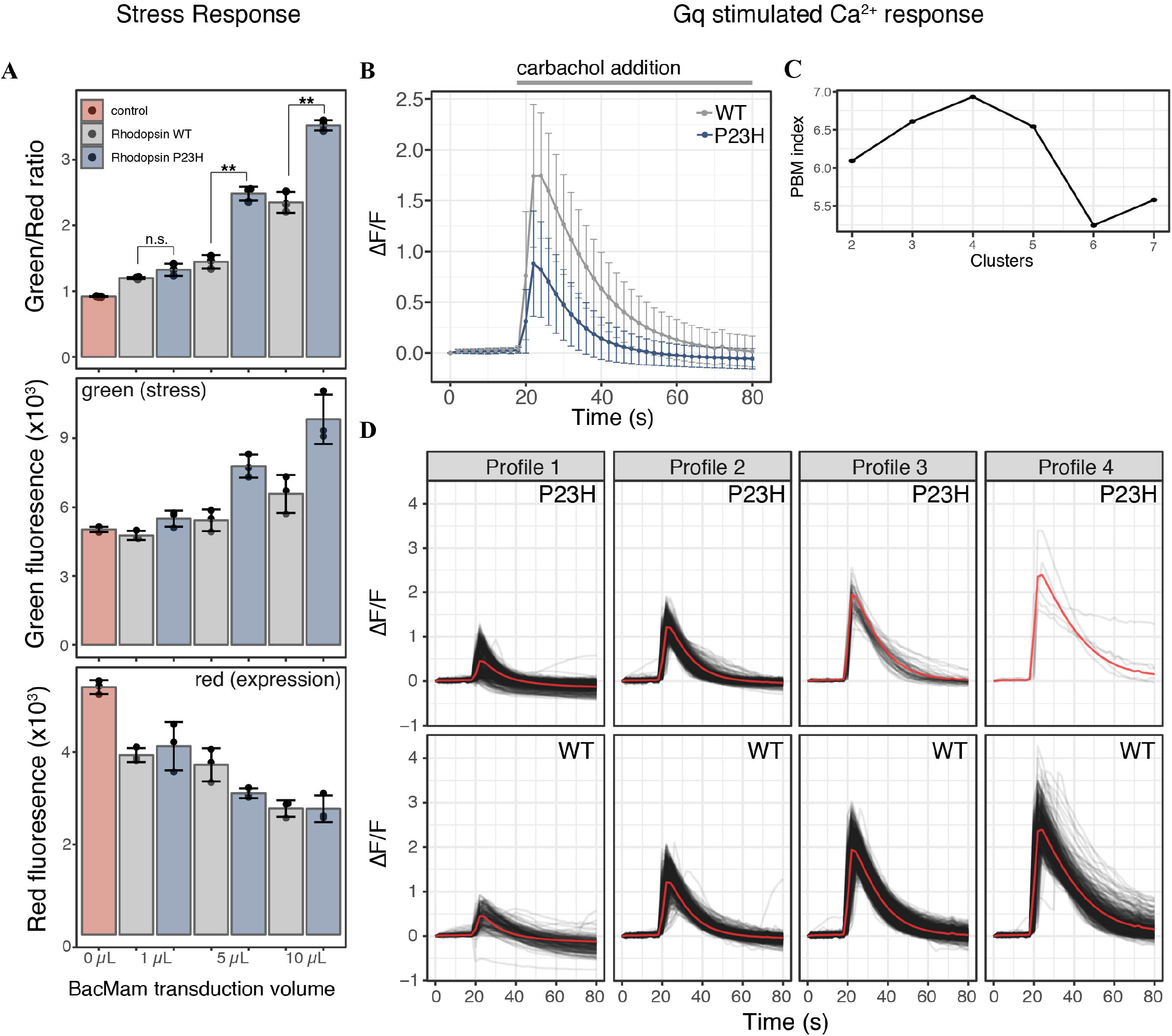
Mutant P23H rhodopsin elicits ER-mediated cell stress and alters Ca^2+^ signaling. **(A)** HEK293T cells transduced with the cell stress biosensor and either no protein (control), WT rhodopsin or P23H rhodopsin. After 24 hours the green, red, and green/red fluorescence ratio were analyzed. Data are plotted as mean ± s.d., n = 3 wells per condition, ** = P-value < 0.01, n.s .= not significant. **(B)** Average profiles of the fold change in cytoplasmic Ca^2+^ in cells expressing the R-GECO Ca^2+^ biosensor and transduced with 5 μL WT or P23H rhodopsin after stimulation with 30 μM carbachol. Data are plotted as the mean ± s.d. of 2 wells and 2125 cells for WT and 2 wells and 1023 cells for P23H. **(C)** PBM index analysis of Ca^2+^ signaling profile clusters in WT and P23H rhodopsin expressing cells. A maximum at 4 clusters indicates Ca^2+^ signaling profiles can be best segmented into 4 distinct profiles. **(D)** Individual Ca^2+^ responses from each cell in (B) were analyzed and separated into four distinct profiles based on the PBM index from (C). Each black line represents a single cell, the red line represents the mean response of each profile.

**Figure 3.**
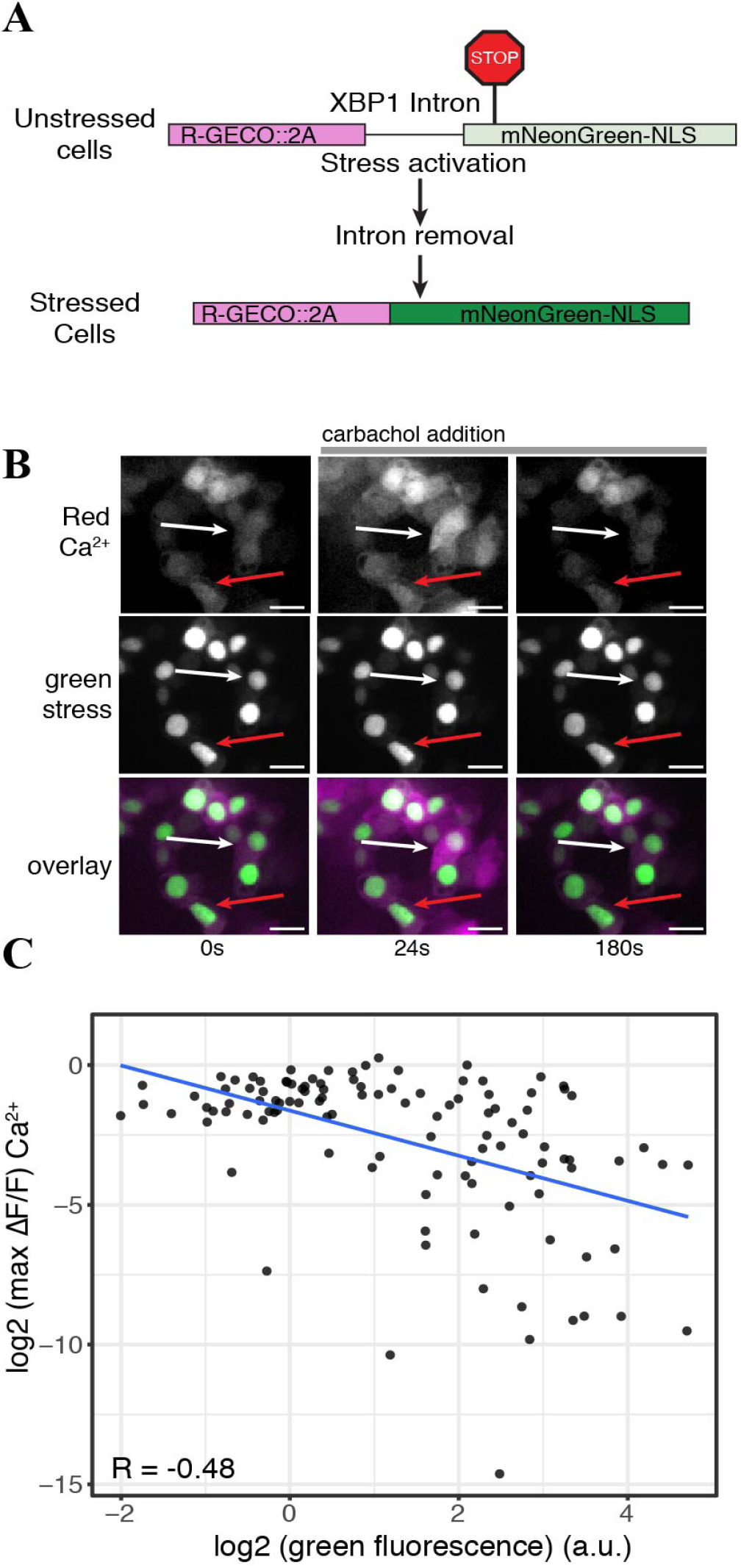
ER stress levels correlate with decreased Ca^2+^ signaling response. **(A)** Schematic of the R-GECO cell stress biosensor. **(B)** Representative image of cells expressing the R-GECO cell stress biosensor and the rhodopsin P23H mutant. Top: red Ca^2+^ channel, middle: green stress channel, bottom: merge. Images are taken before during and after carbachol stimulation. The white arrow depicts a lowly stressed cell. The red arrow depicts a highly stressed cell. Scale bar = 25 μm. **(C)** Scatter plot comparing the stress levels and max fold change in Ca^2+^ levels of individual cells of expressing the R-GECO cell stress biosensor and rhodopsin P23H. The blue line is the linear correlation between the two variables. R = Pearson correlation coefficient.

### Gene Expression

Plasmid transfections were done using Lipofectamine 2000 (Thermo Fisher 11668019) according to the manufacturer’s instructions. Briefly, 0.4 μL of Lipofectamine 2000 per 100 ng of plasmid DNA was added to Opti-Mem reduced serum media (Thermo Fisher 31985088) to create a final volume of 100 μL. After incubation at room temperature for 20 minutes all 100 μL of the transfection mix was mixed with 100 μL of HEK293T cells and plated onto a single well of a 96-well microtiter plate. For all transfection experiments 100 ng of either the cell stress biosensor or the R-GECO-cell stress biosensor plasmids were delivered along with 100 ng of WT or mutant P23H rhodopsin plasmid per 27,000 cells. Four hours later the media was exchanged for 150 μL of Fluorobrite DMEM media. For viral transduction experiments the following BacMam viruses were used: cell stress biosensor (2×1010 viral genes (VG)/mL), R-GECO (3.1×1010 VG/mL), bPAC (1.18×1011 VG/mL), R-cADDis (5.06×1010 VG/mL), rhodopsin, a-synuclein, and TDP-43 viral titers listed in Table 1. For experiments comparing WT and mutant versions of rhodopsin, α-synuclein, and TDP-43 the viral volume was matched to the titer of the WT protein in the case of rhodopsin and TDP-43 and the A53T mutant of α-synuclein. For all experiments using BacMam 25 μL of biosensor was transduced per 48,000 cells. When bPAC was used it was transduced at 5 μL per 48,000 cells. For HEK293T cells all transductions were conducted in the presence of 2 mM sodium butyrate and for SH-SY5Y cells all transductions were conducted in the presence of 6 mM valproic acid. After 24 hours incubation the cells were analyzed for cell stress, Ca^2+^ signaling, or cAMP degradation.

**Table 1.**
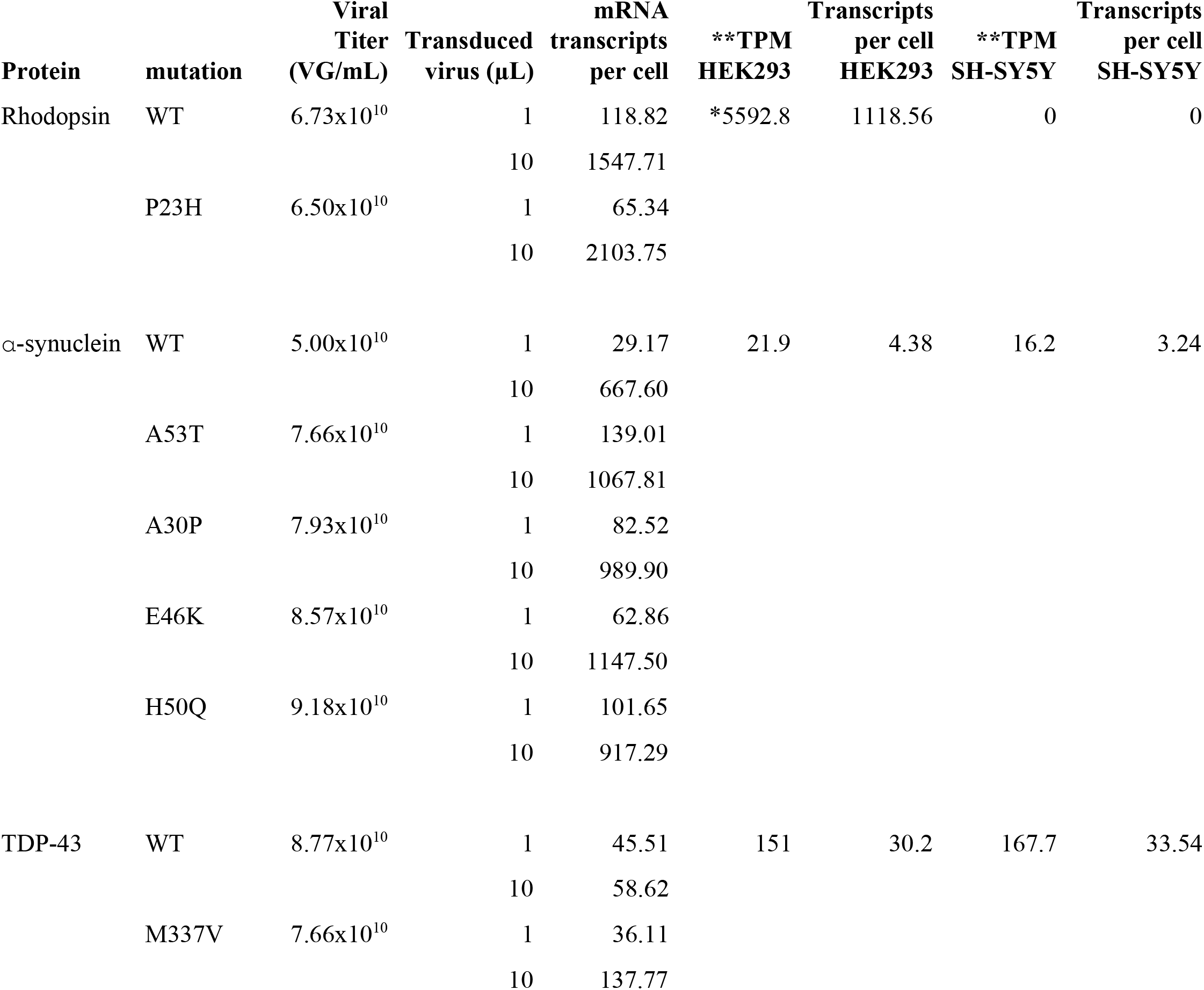
mRNA transcripts per cell for WT and mutant versions of the genes used in this study. All genes were packaged in BacMam and the viral titer was determined as viral genes (VG) per mL. mRNA transcripts per cell were determined by qRT-PCR and compared to endogenous expression levels in HEK293T and SH-SY5Y cells. *For rhodopsin expression data from the retina was used. **Data obtained from (Uhlén et al., 2015), TPM, transcripts per million.

| Protein | mutation | Viral<br>Titer<br>(VG/mL) | Transduced<br>virus (μL) | mRNA<br>transcripts<br>per cell | **TPM<br>HEK293 | Transcripts<br>per cell<br>HEK293 | **TPM<br>SH-SY5Y | Transcripts<br>per cell<br>SH-SY5Y |
| --- | --- | --- | --- | --- | --- | --- | --- | --- |
| Rhodopsin | WT | 6.73x10<sup>10</sup> | 1 | 118.82 | *5592.8 | 1118.56 | 0 | 0 |
|  |  |  | 10 | 1547.71 |  |  |  |  |
|  | P23H | 6.50x10<sup>10</sup> | 1 | 65.34 |  |  |  |  |
|  |  |  | 10 | 2103.75 |  |  |  |  |
| α-synuclein | WT | 5.00x10<sup>10</sup> | 1 | 29.17 | 21.9 | 4.38 | 16.2 | 3.24 |
|  |  |  | 10 | 667.60 |  |  |  |  |
|  | A53T | 7.66x10<sup>10</sup> | 1 | 139.01 |  |  |  |  |
|  |  |  | 10 | 1067.81 |  |  |  |  |
|  | A30P | 7.93x10<sup>10</sup> | 1 | 82.52 |  |  |  |  |
|  |  |  | 10 | 989.90 |  |  |  |  |
|  | E46K | 8.57x10<sup>10</sup> | 1 | 62.86 |  |  |  |  |
|  |  |  | 10 | 1147.50 |  |  |  |  |
|  | H50Q | 9.18x10<sup>10</sup> | 1 | 101.65 |  |  |  |  |
|  |  |  | 10 | 917.29 |  |  |  |  |
| TDP-43 | WT | 8.77x10<sup>10</sup> | 1 | 45.51 | 151 | 30.2 | 167.7 | 33.54 |
|  |  |  | 10 | 58.62 |  |  |  |  |
|  | M337V | 7.66x10<sup>10</sup> | 1 | 36.11 |  |  |  |  |
|  |  |  | 10 | 137.77 |  |  |  |  |

### RT-qPCR

Two wells of a 96-well plate containing 48,000 HEK293T (96,000 total) cells were transduced with either 1 or 10 μL of BacMam virus indicated in Table 1. Cells were then incubated for 24 hours followed by RNA isolation using Quick-RNA Microprep Kit (Zymo Research R1050). cDNA was generated using the M-MLV Reverse Transcriptase kit (Promega M1701) according to the manufacturer’s instructions. qPCR was conducted using Syber select master mix for CFX (Thermo Fisher 4472942).

### Plate reader, Imaging and Image Analysis

For long term plate reader analyses in Figure 1 cells were transfected as described above with the addition of 25 mM HEPES (Thermo Fisher 15630080) to the media. Cells were analyzed for 20-40 hours on a BMG CLARIOstar (Figure 1C) or BioTek SynergyMX (Figure 1D, E) plate reader, heated to 37°C. Reads with excitation/emission of 485/528 and 558/603 with 20 nm bandpass were taken every 30 minutes. High content analysis (Figure 1C and D) was conducted on the Lionheart FX Automated Microscope (BioTek Instruments) at 37°C and 5% CO2. HEK293 cells were transduced with 10 μL of the cell stress biosensor per well in Fluorobrite DMEM containing 6 mM valproic acid and plated onto 96-well plates. 24 hours after plating increasing concentrations of thapsigargin was added to the cells and cells were imaged every hour using a 10x objective and the following filter sets, red fluorescence: 531 Ex, 593 Em LED filter cube, green fluorescence: 469 Ex, 525 Em LED filter cube. Images for Figure 2 were collected on a Zeiss Axiovert 200 using an Olympus UPlanFL 10x/0.30 lens and a Teledyne Qimaging 2000R CCD camera. Images were taken every 2 seconds exciting for 200 ms with a 1A 560 nm LED (ThorLabs DC4100) with a 556/20 Semrock excitation filter and a 617/70 Semrock emission filter. Images for Figure 3 were acquired on an Olympus IX81 using an Olympus UPlanSApo 20x/0.75 lens and a Hamamatsu C9100 EM-CCD camera. Images were acquired every 2s exciting for 200 ms with a 1A Halogen Exfo X-cite series 120 lamp using the same excitation and emission filters. For all bPAC-RcADDis imaging in Figures 4 and 5 the Zeiss Axiovert with the 10x lens was used. Images were acquired every 20 seconds using a 100 ms exposure and the 556/20 Semrock excitation filter. After 100 seconds the sample was pulsed with blue light for 2 seconds using a 1A 470 nm LED and a 470/20 Semrock excitation filter. Following this pulse images were acquired every 20 seconds as before the pulse. All image analysis was conducted using CellProfiler (McQuin et al., 2018). Raw images and Cell-Profiler scripts are available in the supplemental materials.

**Figure 4.**
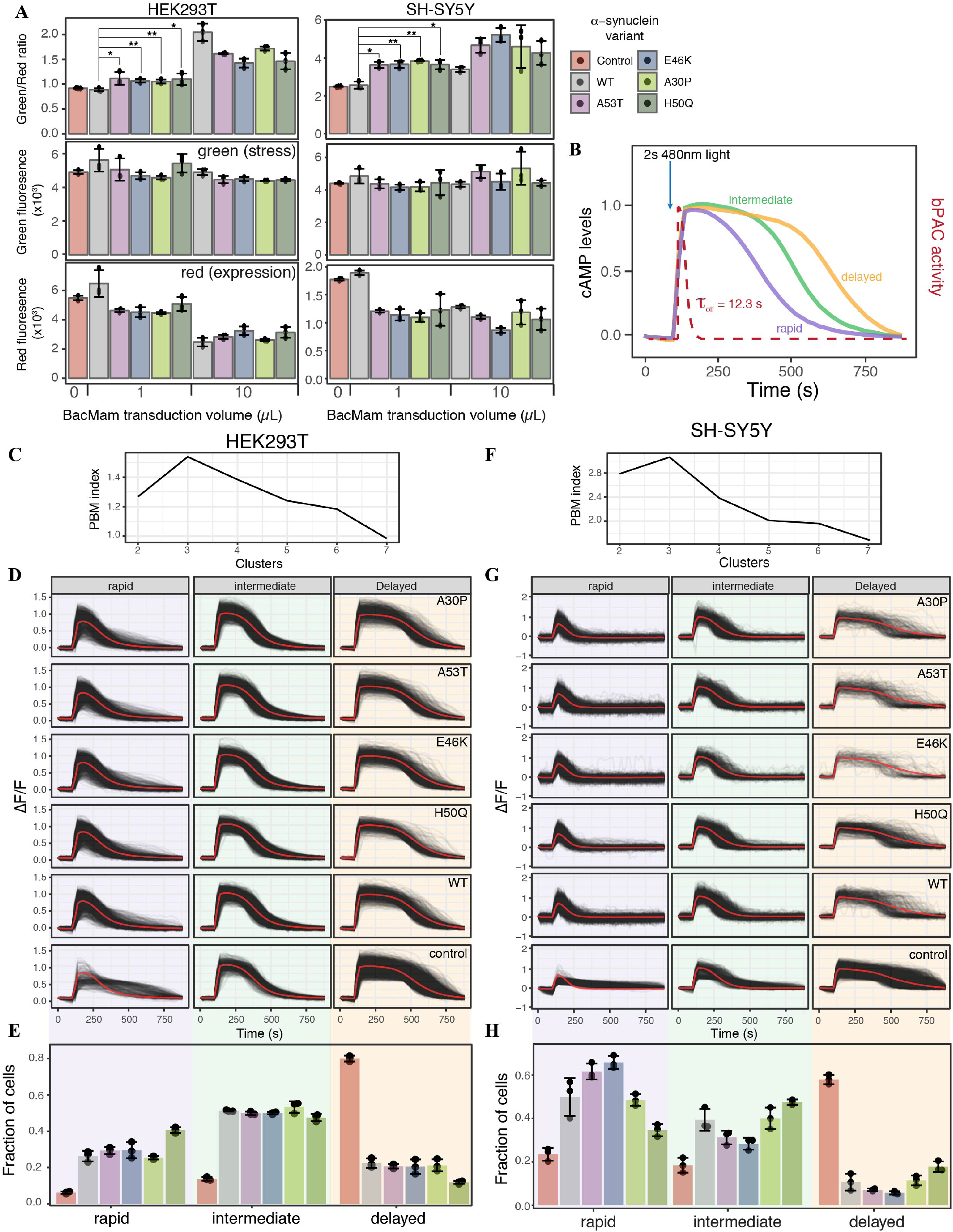
α-synuclein expression induces ER-mediated cell stress and alters cAMP degradation. **(A)** HEK293T or SH-SY5Y cells were transduced with the cell stress biosensor and either no other virus (control) or 1 or 10 μL of WT α-synuclein or one of four α-synuclein mutants. The green, red and green/red ratio fluorescence were measured after 24 hours of expression. Individual data points are plotted and bars represent the mean, error bars represent the standard deviation of n = 3 wells. **(B)** Schematic of PDE activity assay. Cells transduced with the red fluorescent cAMP biosensor R-cADDis and the blue light activated adenylyl cyclase bPAC. Baseline cAMP levels were monitored prior to bPAC activation to raise cAMP levels. A 2 second pulse of blue light activates bPAC to raise cAMP. bPAC activity decays rapidly and cAMP levels were monitored for 15 minutes at which point cAMP levels return to baseline. PDE activity profiles from individual cells revealed three distinct cAMP decay profiles, rapid, intermediate, and delayed. **(C)** PBM index analysis of cAMP degradation profiles in HEK293T cells. Data from control cells or cells expressing WT or a mutant variant of α-synuclein were merged for analysis. A maximum at 3 clusters indicates cAMP degradation profiles can be best segmented into 3 distinct profiles. **(D)** Individual cAMP degradation profiles from control cells and cells expressing WT or a mutant variant of α-synuclein. Each black line represents a single cell, the red line represents the mean response of each profile. N = 3 wells per sample, control = 4047 cells, WT = 3882 cells, A53T = 3385 cells, E46K = 3431 cells, A30P = 3700 cells, H50Q = 3455 cells. (E) Bar plot of the fraction of cells in each profile for control and α-synuclein variants from the profiles determined in (C). N = 3 wells for each condition. Individual data points are plotted along with the mean bar and error bars representing the standard deviation. **(F)**, **(G)** and **(H)** same as in (C), (D), and (E) but for SH-SY5Y cells. N = 3 wells per sample, control = 2697 cells, WT = 2427 cells, A53T = 2775, E46K = 1462 cells, A30P = 1902 cells, H50Q = 2209 cells. ** = P-value < 0.01, * = P-value < 0.05.

**Figure 5.**
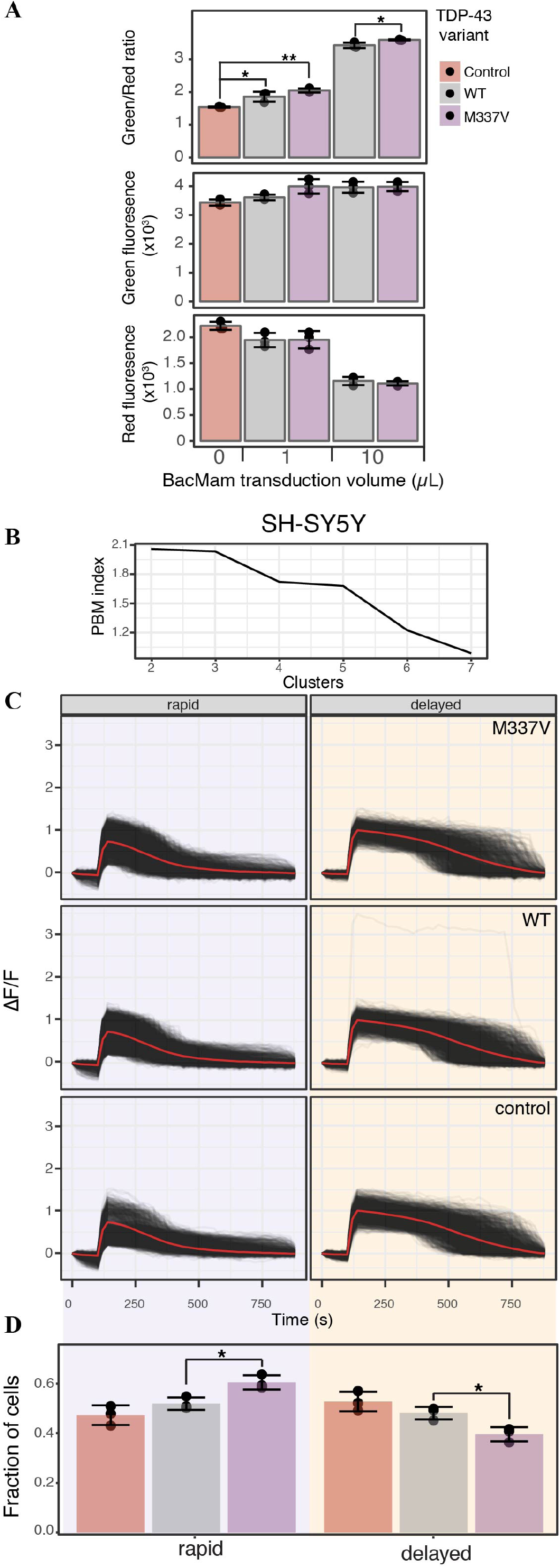
TDP-43 expression induces cell stress and increases cAMP degradation rate. **(A)** SH-SY5Y cells were transduced with the cell stress biosensor and either no other virus (control) or 1 or 10 μL of WT or M337V TDP-43. The green, red and green/red ratio fluorescence were measured after 24 hours of expression. Individual data points are plotted and bars represent the mean, error bars represent the standard deviation of n = 3 wells. **(B)** PBM index analysis of cAMP degradation profiles in SH-SY5Y cells. Data from control cells or cells expressing WT or M337V versions of TDP-43 were merged for analysis. A maximum at 3 clusters indicates cAMP degradation profiles can be best segmented into 3 distinct profiles. **(C)** Individual cAMP degradation profiles from control cells and cells expressing WT or M337V TDP-43. Each black line represents a single cell, the red line represents the mean response of each profile. N = 3 wells per sample, control = 2697 cells, WT = 3352 cells, M337V = 3023 cells. **(D)** Bar plot of the fraction of cells in each profile for control and TDP-43 variants from the profiles determined in (B). N = 3 wells for each condition. Individual data points are plotted along with the mean bar and error bars representing the standard deviation. *= P-value < 0.05.

### Statistical Analysis

Where applicable data are reported as mean ± s.d. of at least 3 replicates. Either the fold change (ΔF/F) in fluorescence or the percent stressed cells was used to determine cell stress response. ΔF/F refers to the fold change in green fluorescence induced by activation of the stress sensor. Percent stressed cells is calculated by dividing the number of cells that display both green and red fluorescence by the total number of cells expressing red fluorescence within a single image. The percent stressed cells analysis was used to analyze high content data in Figure 1. For image analysis in Figures 2-5 individual cells from 2 or 3 images were analyzed, with the total number of wells and cells listed for each experiment in the figure legends. P-values were determined using two-tailed equal variance t-tests. Ca^2+^ signaling and cAMP degradation profile analysis was conducted in R (www.r-project.org) using the TSrepr package (Laurinec, 2018). Data were first filtered for a positive increase in cAMP levels after bPAC stimulation by removing cells displaying a ΔF/F of less than 0.2. The number of cAMP degradation profiles was determined using the PBM index (Pakhira et al., 2004), identifying the number of clusters that had highest PBM index score.

## RESULTS

### A genetically-encoded reversible biosensor to detect cell stress

To assay cellular stress responses we created a genetically encoded live-cell biosensor to detect an ER-mediated stress response through the IRE1α-XBP1 arm of the UPR. Versions of this type of biosensor have been previously described (Iwawaki et al., 2004; Roy et al., 2017), however we sought to modify the sensor to adapt it for use in both imaging and plate reader based assays. Upon detection of misfolded proteins within the lumen of the ER, IRE1α is activated and carries out an unconventional cytoplasmic splicing of the XBP1 transcript. This splicing leads to a frame shift in the XBP1 open reading frame, creating a functional transcription factor which in turn activates a host of stress response genes (Grootjans et al., 2016). Similar to previous versions of this biosensor we co-opted this splicing event to shift into frame a bright green fluorescent protein, mNeonGreen (Figure 1A). We also added a constitutively expressed red fluorescent protein upstream of the XBP1 intron to identify cells expressing the biosensor and as an indicator for protein expression levels. A self-cleaving 2A peptide was placed between the red and green fluorescent proteins to uncouple changes in protein expression from changes in ER-mediated cell stress. Next, we targeted both fluorescent signals to the nucleus for easy image analysis and signal comparison. Lastly, we modified the sensor to better mimic the endogenous UPR. Once the stress has been alleviated in a typical IRE1α-XBP1 response, the spliced transcript and functional transcription factor are degraded (Uemura et al., 2013). Thus, to mimic the endogenous IRE1α-XBP1 response we ensured that the cell stress biosensor was reversible, able to indicate stress activation and be degraded upon stress alleviation. We tested this reversibility by comparing the endogenous XBP1 splicing status to the biosensor splicing status during treatment with thapsigargin, a SERCA pump inhibitor that activates the IRE1α-XBP1 pathway. The splicing of the biosensor and the endogenous XBP1 transcript display similar profiles during a 24 hour treatment with thapsigargin (Figure 1B). The highest level of spliced endogenous XBP1 transcript is observed between two and six hours after thapsigargin treatment, but diminishes within 24 hours (van Schadewijk et al., 2012) (Figure 1B). Both the splicing of the biosensor transcript and the stress related green fluorescence peak between 2-6 hours of treatment, diminishing after 22-24 hours (Figure 1B-D), mimicking the endogenous stress response. Importantly, the cell stress response to thapsigargin is similar whether the stress levels are monitored using the fold change in green fluorescence on a plate reader (Figure 1C, top) or a high content imager using the percent of stressed cells within the well (Figure 1C, middle). Additionally, high content image analysis demonstrates that changes in cell stress occur an order of magnitude prior to changes cell growth (Figure 1C, middle and bottom) when cells are stressed with thapsigargin. These data indicate that the cell stress biosensor may be more sensitive to detect changes in cellular stress response than assays monitoring cell growth or death.

An important feature of many neurodegenerative diseases is sustained cellular stress often caused by genetic mutations or changes in protein expression (Bosco et al., 2011; Ferrari et al., 2011). We next tested the ability of the cell stress biosensor to detect stress mediated through genetic mutations. We chose to assess a wild type and mutant variant of rhodopsin. Rhodopsin is trafficked through the ER to the cell membrane in retinal cells (Deretic and Papermaster, 1991). Mutations in rhodopsin can lead to misfolding of the protein within the ER and are known to cause the progressive blinding disease, Retinitis Pigmentosa (Sung et al., 1991). By co-expressing the cell stress biosensor with either wild type (WT) or a mutant form of rhodopsin known to cause Retinitis Pigmentosa, rhodopsin P23H, the effects of the WT and mutant rhodopsin were monitored over 20 hours. Rhodopsin P23H not only displayed an increased cell stress response (Figure 1E), it also resulted in decreased protein expression (Figure 1E, red). This general decrease in protein expression is a hallmark of the UPR, and coupled with an increase in ER-mediated cell stress demonstrates the ability of the cell stress biosensor to detect cell stress induced by genetic mutations. Lastly, as the cell stress biosensor may be a useful tool for assaying compounds that alleviate the cell stress response, we assessed the ability of the IRE1α inhibitor, 4μ8C to reduce the stress caused by the P23H mutation. After 20 hours of expression, increasing doses of 4μ8C were added to cells expressing the P23H mutation and the subsequent changes in stress related green fluorescence and expression related red fluorescence were monitored. As expected addition of 4μ8C reduced the stress response within five hours after treatment, while no change in general protein expression was observed (Figure 1F). Importantly, 1 μM of 4μ8C, which has an IC50 for IRE1α of 6.8 μM, was able to reduce stress levels of the P23H mutant back to those of WT within 20 hours of treatment (Figure 1F, WT DMSO). These results demonstrate the ability of the cell stress biosensor to detect real-time changes in cell stress responses mediated by genetic components.

### Ca^2+^ signaling is affected by ER stress

Cellular stress responses, including those brought on by ER stress, can lead to a number of changes in cellular physiology. Second messenger signaling is especially sensitive to changes in cell state, thus we sought to explore how ER mediated cell stress, brought on by the overexpression of WT and mutant P23H rhodopsin, affected intracellular Ca^2+^ signaling. We chose to explore Ca^2+^ signaling as the ER is the main store of Ca^2+^ within the cell, and changes to cytosolic Ca^2+^ has been associated with the P23H mutation (Shinde et al., 2016). We first created a modified baculovirus, BacMam, to express the cell stress biosensor along with the WT or mutant forms of rhodopsin. We then transduced HEK293T cells with increasing amounts of WT or mutant rhodopsin BacMam along with BacMam carrying the cell stress biosensor. Analysis of the stress signal, the expression signal, and the green/red ratiometric signal (Figure 2A) demonstrated a dose dependent increase in cell stress mediated by WT and mutant P23H rhodopsin, when compared to control HEK293T cells that were not transduced with either rhodopsin construct. The stress response from P23H rhodopsin became distinguishable from the WT stress response at 5 μl transduced virus, which corresponds to a similar level of rhodopsin mRNA expression observed in the retina (Table 1, (Uhlén et al., 2015)). To determine if the stress response instigated by the P23H mutation affected Ca^2+^ signaling we co-transduced 5 uL of either WT or P23H rhodopsin BacMam along with the R-GECO Ca^2+^ biosensor (Wu et al., 2013) and the human cholinergic receptor muscarinic 1 (hM1) GPCR. Cells expressing either WT or P23H rhodopsin were then stimulated with carbachol to activate a Gq signaling response. Cells were imaged every two seconds before and during carbachol treatment. As shown in Figure 2B the average Ca^2+^ response was quite different between the cells expressing WT and P23H rhodopsin. WT cells displayed an average ~1.75 fold change in cytoplasmic Ca^2+^ levels whereas P23H cells displayed only an average ~0.8 fold change. Inspection of individual Ca^2+^ traces from single cells imaged in Figure 2B suggested there may be distinct subpopulations of cells with variable responses to Gq activation. To explore this possibility, single cell traces for the WT and P23H data were analyzed for Ca^2+^ response. All single cell traces from both the WT and P23H expressing cells were pooled and analyzed for the presence of unique Ca^2+^ signaling patterns. To identify the number of distinct Ca^2+^ signaling clusters present in the data we tested the goodness of fit for 2-7 clusters of Ca^2+^ signaling profiles. Use of the PBM index (Pakhira et al., 2004) to validate cluster number, we identified four Ca^2+^ response profiles (Figure 2C). Two profiles, dominant in the WT cells, displayed increased Ca^2+^ responses upon hM1 activation (Figure 2D, profiles 3 and 4), resembling a typical Gq response (Tewson et al., 2013). The other two profiles, dominant in the P23H cells, displayed a blunted Ca^2+^ response (Figure 2D, profiles 1 and 2). However, the presence of profiles 1 and 2 in cells expressing WT rhodopsin suggests that even overexpression of WT rhodopsin elicits altered Ca^2+^ signaling along with increased ER stress (Figure 2A, D).

Analysis of ER-mediated cell stress and individual Ca^2+^ profiles suggests variability in Ca^2+^ signaling response may be related to varying levels of cellular stress induced by the rhodopsin P23H mutation. To explore this possibility we created a modified version of the cell stress biosensor where the constitutively expressed red fluorescent protein was replaced with R-GECO (Figure 3A). We then co-expressed this version of the biosensor with rhodopsin P23H and hM1 and again activated Gq signaling through carbachol treatment. As seen in Figure 3B, cells with higher levels of cell stress had blunted Ca^2+^ responses, while those with lower levels of cell stress displayed a more typical Gq Ca^2+^ response. Comparing the fold change in Ca^2+^ levels upon Gq activation with stress levels revealed a negative correlation between cell stress and Ca^2+^ signaling. The higher the levels of cell stress, the more blunted the Gq Ca^2+^ response (Figure 3C). Together these data demonstrate that not only are Ca^2+^ signaling dynamics dependent upon cell stress, but that the degree of signaling disruption is dependent upon the degree of cell stress.

### α-synuclein overexpression induces cell stress and alters PDE activity

ER stress is also associated with protein folding diseases of proteins that are not directly trafficked through the ER (Scheper and Hoozemans, 2015). To test the ability of the cell stress biosensor to detect indirect ER stress we created BacMam constructs of wild type α-synuclein (α-syn) and four α-syn mutants associated with Parkinson’s Disease (Maiti et al., 2017). HEK293T and SH-SY5Y cells were co-transduced with BacMam containing the cell stress biosensor and either 1 μL or 10 μL of BacMam containing either WT or one of the mutant α-syn genes. SH-SY5Y cells were chosen along with HEK293T cells as they are a neuroblastoma cell line that has become an increasingly useful neuronal model to study not only Parkinson’s disease but other neurodegenerative diseases as well (Nonaka et al., 2009; Vasquez et al., 2018; Xicoy et al., 2017). After 24 hours of expression both the green fluorescence levels, to monitor stress induction, and the red fluorescence levels, to monitor protein expression, were analyzed using a plate reader. In both HEK293T and SH-SY5Y cells all of the α-syn mutants displayed increased ER-mediated cellular stress, compared to the WT at both 1 μL and 10 μL of virus (Figure 4A, green/red ratio). Notably, in both cell lines the α-syn mutants induced a significant decrease in overall protein expression at 1 μL of virus, which is equal to ~7x overexpression (Figure 4A red, Table 1). Normalizing the green stress fluorescent signal to this overall drop in expression results in significant detection of cell stress in each of the α-syn mutants. Interestingly, in HEK293T cells at 10 μL of virus the WT α-syn induces increased stress levels compared to the mutants, but in SH-SY5Y cells each α-syn mutant induced greater stress levels than the WT. This result suggests cell type may be an important factor to consider when assessing the effects of α-syn mutations on cellular responses. Together, these data demonstrate the ability of the cell stress biosensor to detect ER-mediated cell stress induced by genetic mutations affecting proteins in the cytoplasm as well as the ER.

Recent studies have identified phosphodiesterase (PDE) inhibitors as a way to preserve dopaminergic neurons (Morales-Garcia et al., 2011), promote neurogenesis (Morales-Garcia et al., 2015) and rescue Parkinsonian phenotypes (Bartolome et al., 2018). PDE inhibitor treatment increases cAMP levels within the cells which leads to increased CREB activation and expression of genes promoting neurogenesis (Morales-Garcia et al., 2011). However, how the regulation of basal levels of cAMP is altered in the disease state remains unclear. We reasoned that in cells expressing variants of α-syn changes to PDE activity, either through changes in expression or enzyme activity, may alter cAMP degradation kinetics leading to altered cAMP regulation. Thus, we assessed the effects of expressing different α-syn variants on cAMP levels and PDE activity in HEK293T and SH-SY5Y cells.

To remove the need for receptor activation to stimulate cAMP production we developed a completely optical based approach to monitor cAMP production and degradation. First, we used a blue light-activated adenylyl cyclase, bPAC (Stierl et al., 2011) to transiently raise cAMP levels within the cell, bypassing the endogenous receptor and adenylyl cyclase. The rapid off rate of bPAC allows direct monitoring of cAMP degradation by PDEs using R-cADDis, a red cAMP biosensor (Figure 4B). We first compared the PDE activity profiles between control cells and cells transduced with 1 μL of either WT or mutant α-syn, the lowest level of overexpression found to increase cellular stress. We again used the PBM index to determine the number of distinct cAMP degradation profiles present within the different control and α-syn expressing cells (Figure 4C and F). In both HEK293T and SH-SY5Y cells three unique cAMP degradation profiles were observed, rapid, intermediate, and delayed (Figure 4D and G). Overexpression of either WT or mutant forms of α-syn resulted in drastic changes in the cAMP degradation profile distribution in both cell types. In HEK293T cells overexpression of any form of α-syn displayed a shift in cAMP degradation away from the delayed profile towards the rapid and intermediate profiles (Figure 4E). Similar shifts towards more rapid cAMP degradation were observed in SH-SY5Y cells as well. However, in these cells the A53T and E46K mutations displayed an even greater shift towards the rapid profile when compared to WT α-syn (Figure 4H). The fact that overexpression of WT α-syn creates a shift towards increased PDE activity, with no observable change in ER-mediated cell stress, suggests that monitoring changes in cAMP degradation may be a more sensitive readout to detect defects in cellular function than ER-mediated cell stress. Or that changes to PDE activity may precede cell stress in overexpression models of WT α-syn. Notably, overexpression of WT α-syn has been shown to recapitulate Parkinson’s symptoms (Chesselet, 2008; Mochizuki et al., 2006). Together, these data suggest that in the presence of minimally overexpressed α-syn variants, cells undergo an ER-mediated cellular stress response and display an increased PDE activity. These results are consistent with previous reports that PDE inhibitors can preserve dopaminergic neurons suggesting that cAMP levels play an important role in the progression of Parkinson’s disease (Morales-Garcia et al., 2011, 2015).

### TDP-43 overexpression increases cAMP degradation rate

To determine if these changes in cell stress and cAMP degradation kinetics are unique to α-syn, we repeated the cell stress assay and optical interrogation of PDE activity using SH-SY5Y cells co-transduced with either WT or the M337V mutant of TDP-43. Overexpression of either WT or M3337V versions of TDP-43 resulted in increased cell stress (Figure 5A), when compared to control cells at 1 μL of transduced virus, which corresponds to ~1.5x overexpression of TDP-43 in SH-SY5Y cells (Table 1). Transduction of 5 μL of virus, corresponding to ~5x overexpression, resulted in increased cell stress when compared to control cells. Further, when compared to WT, the M337V mutation displayed a significant increase in cell stress (Figure 5A, 10 μL). Analysis of cAMP degradation profiles by PBM clustering revealed two distinct cAMP degradation patterns, a rapid and a delayed profile (Figure 5B). As control cells are analyzed along with cells expressing TDP-43 WT or TDP-43 M337V to determine cluster number the intermediate profiles identified when comparing control cells to α-syn variants were segmented into either rapid or delayed profiles. Similarly to α-syn overexpression, both the WT and M337V mutant of TDP-43 displayed a shift towards rapid cAMP degradation profiles at 1 μL of virus (Figure 5C and D). The M337V mutant also displayed a significant shift away from the delayed cAMP degradation profile towards the rapid profile when compared to WT TDP-43 (Figure 5D). These data are not only consistent with overexpression of TDP-43 increasing PDE activity, but with the notion that detectable changes in cell signaling are observed prior to changes in ER-mediated cell stress when comparing WT and M337V TDP-43 variants. These data, coupled with the similar results for α-syn, suggest that activation of cell stress and increased PDE activity are a common feature of overexpression of two of the major proteins associated with Parkinson’s Disease and ALS.

## DISCUSSION

The cell stress biosensor described here builds upon previous attempts to create ER stress biosensors through the IRE1α-XBP1 pathway (Iwawaki et al., 2004; Roy et al., 2017). The generation of a reversible ER stress biosensor makes it possible to identify the onset of cell stress, and to search for new drugs that will alleviate this stress. This feature allows for screening of compounds or genetic mutations that activate ER stress as well those that inhibit the response, using a single assay. The detection of ER stress brought on by a mutation in rhodopsin, and the subsequent inhibition of this stress response by blocking IRE1α splicing activity through the use of 4μ8C is an intriguing example of such an intervention. Moreover, these assays were conducted in HEK293T cells, but recapitulated the stress response observed in retinal cells. This This type of assay, where both activation and inhibition of a cellular stress response can be detected in an immortalized cell model should prove useful.

Diseases such as Retinitis Pigmentosa, Parkinson’s Disease and ALS are difficult to study and treat because the assays used to study the effects of these diseases are often endpoint assays monitoring cell death. At this point the ability to intervene and alter the course of the affected cell is lost. Furthermore, any changes in cell function, such as susceptibility to neurotransmitters and agonists, changes to second messenger regulation, or organelle function are best studied during the disease process. Many neurodegenerative diseases not only display cellular stress responses but changes in cellular signalling events as well (Shinde et al., 2016; Xu et al., 2012). The changes observed in cAMP and Ca^2+^ signaling induced by neurodegenerative mutations observed here also raises the question of how affected cells respond to stimuli. Changing the balance and timing of second messenger signaling in the cell may affect the ability of the cell to properly respond to, or transduce, extracellular signals such as neurotransmitters and neuromodulators. This alteration in cellular transduction status may also impact the druggability of cells. For example, a neuron expressing the A53T α-syn mutation displays an increased cell stress response and altered cAMP regulation. However, a neighboring cell, where the mutation has not caused build up of misfolded protein yet, has intact cAMP regulation. If both cells are treated with agonists to the dopamine receptor there will likely be unique signal transduction responses in each cell. These differences are likely to affect how the cell responds or does not respond to the stimuli. These changes are important to consider in the development of drugs targeting these diseased cells. If a cell has lost its ability to transduce certain signals, those receptors, even though present at the cell surface may no longer be a valid drug target. Indeed, our results suggest that targeting PDE activity may be a more effective method to restore the cellular transduction status. Thus, understanding of how multiple stress and signaling pathways merge to affect cellular function in neurodegeneration will be an important step in the search for drugs to combat these diseases.

Finally, while ER-mediated cell stress is an important stress response involved in neurodegenerative disorders many other cell stress pathways are involved. Proteasome stress, protein aggregation, mitochondrial stress, and oxidative stress have all been associated with neurodegenerative disease (Ciechanover and Kwon, 2017; Tan et al., 2019). Creating and combining genetically-encoded fluorescent biosensors for these pathways with biosensors for cellular signaling would further enhance our understanding of how neurodegenerative diseases disrupt cellular homeostasis and create new avenues for treatment of these diseases.

## ACKNOWLEDGEMENTS

This work was supported by the ASEE/NSF Small Business Postdoctoral Research Diversity Fellowship Program, NSF award IIP-1552305 and The National Institute of Neurological Disorders and Stroke, Small Business Innovation Research grant 1R43NS108817-01. We thank members of Montana Molecular for their input on the project.

## AUTHOR CONTRIBUTIONS

K.M.H. and T.E.H. designed the experiments, K.M.H. directed the project. K.M.H. and T.E.H. wrote the manuscript. E. C.R. conducted RT-qPCR, RT-PCR, viral transductions and cell culture. S.M. produced BacMam virus. K.M.H. conducted cell culture, transfections and transductions, collected imaging data, plate reader data, and analyzed the data. J.E.C. conducted high content experiments and analysis.

## SUPPLEMENTARY MATERIAL

All raw images and Cell Profiler analysis scripts are available on figshare at https://doi.org/10.6084/m9.figshare.9209486.v1

